# Rapid Raman spectroscopy-based test for antimicrobial resistance

**DOI:** 10.1101/2024.08.07.606953

**Authors:** Vladimir Mushenkov, Ksenia Zhigalova, Pavel Denisov, Alexey Gordeev, Dmitry Lukyanov, Vladimir Kukushkin, Tatiana Priputnevich, Elena Zavyalova

**Affiliations:** Chemistry Department of Lomonosov Moscow State University, Moscow, Russian Federation; National Medical Research Center Obsterics, Gynecology and Perinatology the name of Academician V.I. Kulakov, Moscow, Russian Federation; Center for Molecular and Cellular Biology, Skolkovo Institute of Science and Technology, Moscow, Russian Federation; Osipyan Institute of Solid State Physics Russian Academy of Science, Chernogolovka, Russian Federation

**Keywords:** antibiotics, antimicrobial resistance, MTT, Raman spectroscopy

## Abstract

Antimicrobial resistance is one of the top global health threats. In 2019, antimicrobial resistance was associated with 4.95 million deaths, of which 1.97 million were caused by drug resistant infections directly. The main subset of AMR is the antibiotic resistance, that is resistance of bacteria to antibiotic treatment. Traditional and most commonly used antibiotic susceptibility tests are based on detection of bacterial growth and its inhibition in the presence of an antimicrobial. These tests typically take over 1-2 days to perform, so empirical therapy schemes are often administered before the proper testing. Rapid tests for antimicrobial resistance are necessary to optimize the treatment of bacterial infection. Here we combine MTT test with Raman spectroscopy to provide 1.5-hour long test for minimal inhibitory concentrations determination. Several *E*.*coli* and *K*.*pneumoniae* strains were tested with three types antibiotics, including ampicillin from penicillin family, kanamycin from aminoglycoside family and levofloxacin from fluoroquinolone family. The test provided the same minimal inhibitory concentrations as traditional Etest confirming its robustness.

## Introduction

Antimicrobial resistance (AMR) is one of the top global health threats. In 2019, antimicrobial resistance was associated with 4.95 million deaths, of which 1.97 million were caused by drug resistant infections directly^1^. The main subset of AMR is the antibiotic resistance, that is resistance of bacteria to antibiotic treatment. Antibiotic resistance threat escalates quickly with widespread, and often incorrect, use of antibiotics^2^. The significant decline in new antibiotic discoveries exacerbates the problem even more. Most antibiotics were discovered in 1950-1970s, in the “Golden Age of antibiotics”, and since 1987 no new class of antibiotics has been discovered for use as treatment^3,4^.

In United States alone, antimicrobial resistance increased healthcare cost by $20 billion, and it is estimated to cause $35 billion loss in productivity annually^5^. Drug resistant infections are difficult to treat, resulting in more expensive therapy and longer hospital stay. Even if therapy is administered, there is significant rate of its inefficiency, with therapy failure rates as high as 54% for MRSA (Methicillin-resistant Staphylococcus aureus) infections^6^.

In drug resistant infections treatment, fast therapy appointment is crucial. For life – threatening conditions, such as sepsis, appropriate therapy needs to be applied within the first hours after diagnosis^7^. Traditional and most commonly used antibiotic susceptibility tests (ASTs) are based on detection of bacterial growth and its inhibition in the presence of an antimicrobial^8^. These ASTs typically take over 1-2 days to perform, so empirical therapy schemes are often administered before the results of AST^9^. Because of the delay, AST results are often not used at all^10^. The delayed appropriate antibiotic therapy can result in prolonged hospital length of stay and higher mortality rate^11^.

### MTT assay for assessing bacterial cell viability

MTT assay is based on the enzymatic reduction of tetrazolium salts, commonly MTT (3-(4,5-dimethylthiazol-2-yl)-2,5-diphenyltetrazolium bromide) to violet insoluble formazan (Figure 1)^12^. In living eukaryotic cells, MTT can be reduced by oxidoreductase and dehydrogenase enzymes, mainly associated with electron transport chain in mitochondria^12^. The reaction rate depends on cell metabolic activity; therefore, only living and active cells will sustain efficient MTT reduction. The amount of formazan formed correlates with the cell metabolism level and their viability in the drug-containing media ^13^. MTT reaction is applied for cell proliferation estimation^14,15^, cytotoxicity evaluation^16,17^. Various anti-cancer drug screenings rely on the MTT test to detect tumor cell suppression^18,19^.

**Figure 1.**
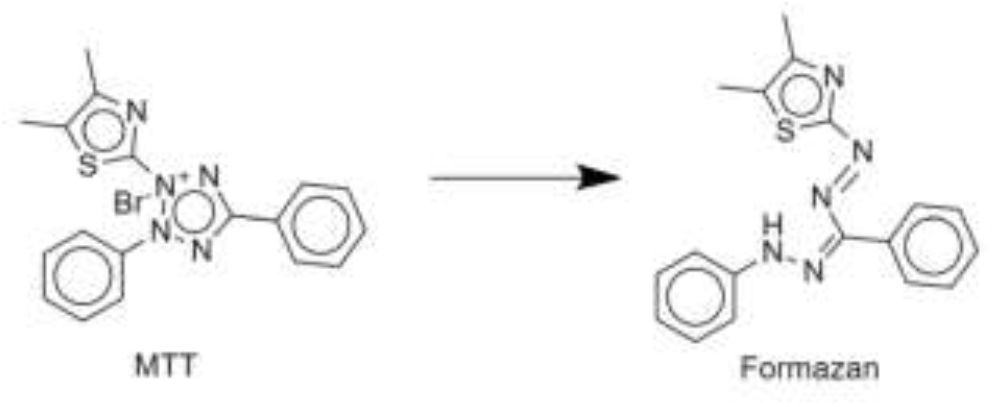
MTT reaction. MTT reagent reduction by cellular reductase enzymes produces a colored product, formazan.

While MTT assay was originally designed as an assay for eukaryotic cells, this method was also used for microbial cells viability estimation^20^. Although the mechanism of MTT reduction in bacteria is not understood well, there are numerous reports which apply MTT assay for bacteria, either gram-positive or gram-negative, including multidrug-resistance strain identification ^21,22^, antibiotic minimal inhibitory concentration (MIC) evaluation^23,24^, biofilm formation^25^ and neutrophilic bactericidal activity evaluation^26^. MTT assay was applied for estimating the number of growing cells of *M. tuberculosis* strains for rapid evaluation of drug resistance. In this case, MTT assay allowed to perform test within 5-7 days, instead of 3-4 weeks for standard colony counting method^27^.

### Raman spectroscopy

Raman spectroscopy is a label-free, noninvasive technique, which is based on nonelastic light scattering^28^. Raman scattered light has a shift specific to vibrational states of the molecule, and Raman spectrum is a unique fingerprint of the substance^29^. Relatively low efficiency of inelastic scattering could be a limitation to sensitivity of analyses, so special modification was developed to overcome low light intensity – surface-enhanced Raman spectroscopy (SERS) and resonance Raman spectroscopy (RRS)^30^.

Surface–enhanced Raman spectroscopy intensifies signal due to surface plasmon resonance effect. This effect occurs on rough metal surfaces, such as surfaces of nanoparticles, the most commonly used SERS-active substrate^31^ Signal is enhanced up to 10^14^ times, which brings SERS to single molecule level of detection^32^. However, signal amplification highly depends on the properties of the SERS-active surface; and the production of reproducible substrates is a rather complex task^33^.

Resonance Raman spectroscopy occurs when the excitation light wavelength is close to absorption maximum of a molecule and light frequency could become in resonance to frequency of molecular electronic transitions^34^. In addition to signal enhancing (up to 10^6^ times), in resonance Raman spectrum additional bands are emerging, which refers to transitions undetectable by standard Raman scattering, such as overtones and combinational modes, therefore additional information can be obtained from the spectrum^35^.

### A combination of resonance Raman spectroscopy and MTT assay

Although absorbance spectrophotometry in visible range of the spectrum is a common practice for formazan detection in MTT assay, it has some disadvantages. First of all, formazan absorbance results are sensitive to the media, so the extraction step is necessary. Even after the extraction of formazan from the sample, formazan spectrum could be interfered by protein precipitation caused by organic solvent^36^. Spectrophotometric assay lacks sensitivity taking several hours for the assay.

Raman spectroscopy provides a characteristic spectrum of formazan that can be used for a specific determination of the compound in a complex media. Characteristic formazan spectral lines are not interfered by other substances in the sample, so steps of extraction and purification of formazan are not required. Sensitivity of Raman spectroscopy detection of formazan is comparable with those of spectrophotometry. But the sensitivity of Raman scattering could be increased substantially with use of surface-enhanced or resonance Raman approaches^37,38^

The intensity and the Raman shift of characteristic peaks of formazan and MTT depends on the excitation light wavelength. Optimal wavelengths for MTT and formazan spectra measurements are 532 and 633 nm respectively.633 nm laser provides a resonance Raman spectrum of formazan^39^. Comparable sensitivities for both MTT and formazan could be obtained using excitation wavelength of 532 nm. Several peaks in formazan spectrum were shown to have a linear dependence on formazan concentration being suitable for a quantitative analysis. Namely, a peak at 967 cm^−1^ for 532 nm excitation wavelength and the peaks at 722 cm^−1^ and 967 cm^−1^ for 633 nm excitation wavelength are valuable for a quantitative analysis^40^.

In this study, we developed a rapid AST that is based on a combination of resonance Raman spectroscopy and MTT assay. The method has turnaround time of about 2 hours (disregarding time needed for cell growth for samples with low concentration) having low labor intensity. We used the method on antibiotic resistant and susceptible strains of *E. coli* and *K. pneumoniae* to estimate MICs. The estimated MICs were compared with MICs obtained by a common technique, broth dilution.

## Results and discussion

### Raman spectra of MTT, formazan and bacteria after MTT treatment

In this work the portable Raman spectrometers with lasers of 532 nm and 637 nm wavelengths were used. The spectrometers were equipped with holders for 2 mL sealed vials. The devices allow investigation of liquid biological samples with no cross-contamination inside the spectrometers.

Firstly, we compared the suitability of these spectrometers for formazan detection in the suspension of living bacterial cells. Figure 2 shows the comparison of Raman spectra of MTT and formazan at two excitation wavelengths. MTT and formazan have unique Raman spectra that allows discrimination these two substances in the mixtures. 532 nm and 637 nm excitation wavelengths provided nearly equal intensity for MTT. However, intensities of formazan spectra differed by an order of magnitude being much higher with 637 nm excitation wavelength. This observation is consistent with previous study that reported resonance Raman spectrum of formazan at 633 nm excitation wavelength of Raman spectrometer.^39,40^

**Figure 2.**
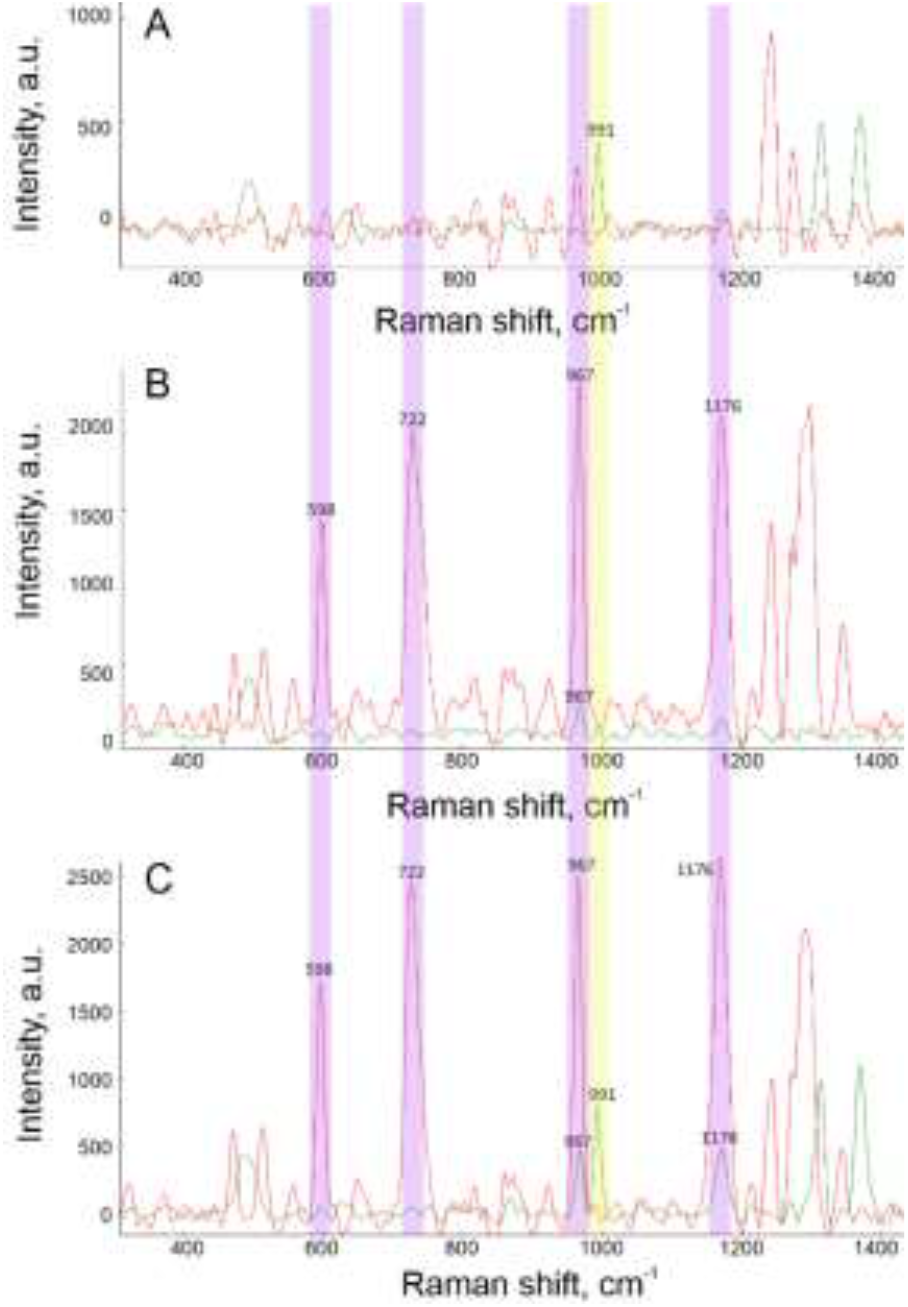
Raman spectra of 0.4 g/L MTT solution in water (A), 0.02 g/L formazan suspension in water (B), E.coli after incubation with MTT reagent. Spectra were collected with 532 nm and 637 nm excitation wavelengths of Raman spectrometer (green and red colors, respectively). Characteristic bands are signed; MTT band is shown in yellow; formazan bands is shown in purple.

Raman spectroscopy allows decoding of a mixture of MTT, formazan and living cells in the cultural medium. The representative spectra are shown in Figure 2C. *E*.*coli* bacterial culture (2 MFU) was incubated with MTT for 30 minutes. Spectra of bacterial culture have characteristic bands for both compounds. Spectra with 532 nm excitation wavelength indicate nearly half-conversion of MTT into formazan. In the case of 637 nm excitation wavelength MTT signal is barely visible, while formazan signal is high due to the resonance effect. Both excitation wavelengths can be used to estimate MTT conversion in formazan; but 637 nm excitation wavelength was considered to provide higher sensitivity due to the enhancement supported by the resonance effect.

### Development of the Raman spectroscopy-based antimicrobial resistance test

Firstly, we have found that night cultures in stationary growth phase provide a poor RRS signal. The cultures were diluted with a fresh medium and incubated for 1 hour at 37°C. The RRS signal increased more than 5 times as a result. Changes in visible range of the spectrum are commonly used to estimate the quantity of bacteria in the cultures.^41,42^ The increase in cell quantity during the elaborated AMR test was assessed with optical density at 600 nm. We found no statistically significant changes in bacterial titer for the samples with 0.5-5 MFU. Possibly, *E*.*coli* were in the lag-phase during 1-2 hours for the rapid test performance.

Next, formazan accumulation was determined with absorbance spectroscopy and Raman spectrometry. The wavelength of 570 nm was used for formazan detection in the visible range. The optical density changes nearly linearly; however, estimation of the samples with low bacterial concentrations (<2 MFU) was not linear. Possible reasons include the increase of light scattering contribution and rather large standard deviations compared to the absolute values (Figure 3A). The Raman spectroscopy provided a smooth monotonous dependence in the whole concentration range studied with a linear range from 2 to 16 MFU (Figure 3B). Raman spectroscopy provided the specific determination of formazan instead of the composite signal in absorbance spectrometry.

**Figure 3.**
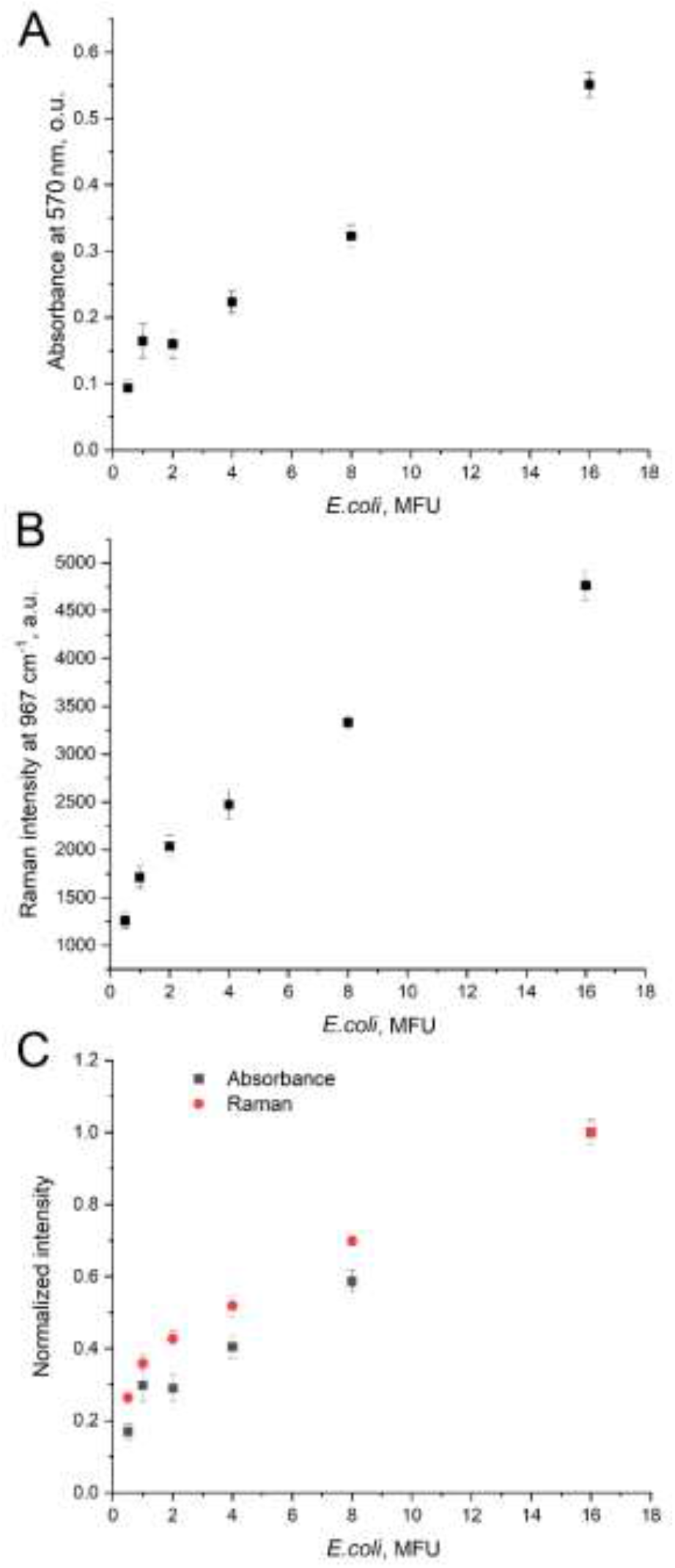
A comparison of optical density of MTT-treated E.coli samples (A) and Raman spectra intensity of MTT-treated E.coli (B) in different bacterial titer. Normalized values were acquired by dividing the values from (A) and (B) by the values for 16 MFU samples (C).

RRS is more precise instrument for low content of bacteria. Normalized spectra (Figure 3C) are rather close to each other. RRS provided nearly the same results as absorbance spectroscopy emphasizing the relationship between the quantity of the bacteria and their enzymatic activity in MTT test.

Next, the antibiotic susceptibility assay was tested. Levofloxacin resistant and susceptible *E*.*coli* strains were treated with 13 µg/mL of levofloxacin for 1 hour at 37°C. Then the MTT reagent was added to the probe, and formazan accumulation in the probe was studied with RRS. The levofloxacin resistant *E*.*coli* strain gave a plateau of formazan RRS intensity after 30 minutes of incubation. Similarly, levofloxacin susceptible *E*.*coli* strain reached plateau of the signal after 30 minutes of incubation; but the intensity of the signal was nearly 2 times lower compared to the resistant strain (Figure 4A). Possibly, the formazan signal was not diminished completely as cells were not killed but exerted toxic effects from the antibiotic. The time of incubation with MTT was fixed at 20 min for the further experiments in order to reach a plateau.

**Figure 4.**
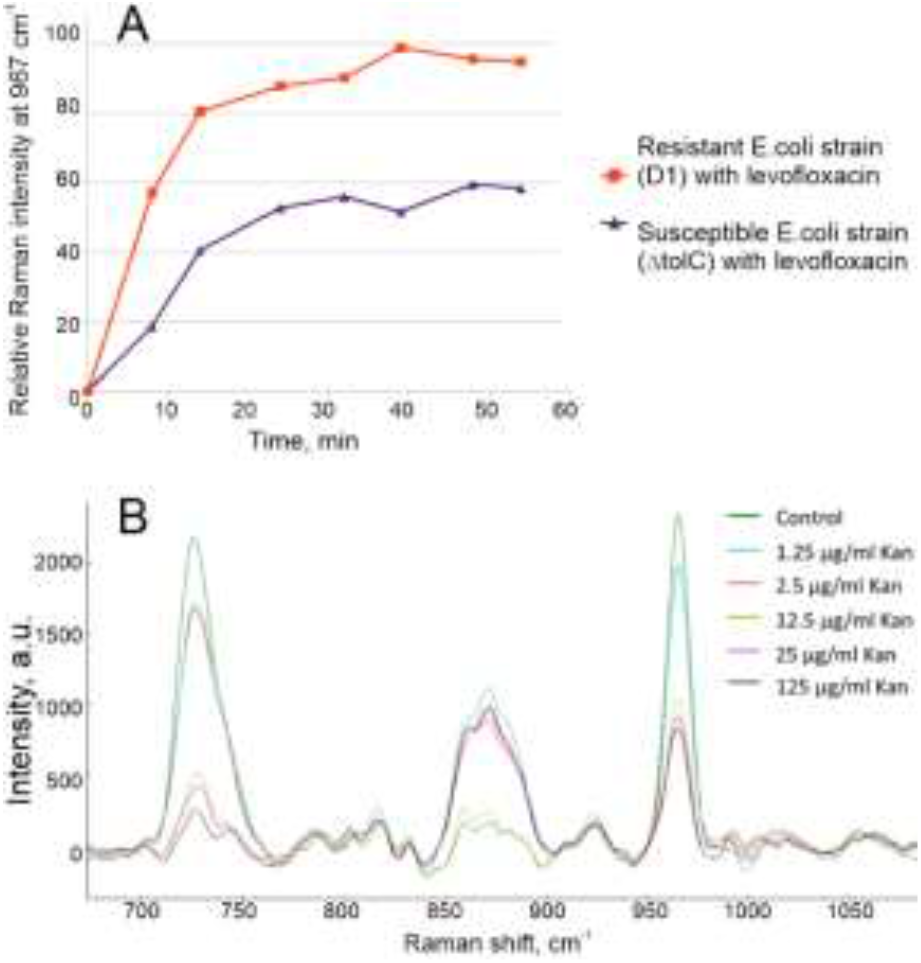
Time dependence of Raman intensity for levofloxacin resistant (D1) and susceptible (ΔtolC) E.coli strains in the presence of 13 µg/mL of levofloxacin (A). Raman spectra of kanamycin susceptible strain of E.coli (D1) after treatment with different concentrations of kanamycin (B).

The concentration dependence for another antibiotic is shown in Figure 4B. *E*.*coli* D1 strain was treated with kanamycin in different concentrations. Increasing concentration of the drug leads to decrease of the formazan signal intensity. The signal changed abruptly from nearly 100% to 50% at 2.5 µg/mL. This value might be due to residual metabolic activity of some enzymes, or by some reducing agents.^43–46^

### Performance of the AMR test on the antibiotic resistant and susceptible *E*.*coli* strains

Two pairs of antibiotic susceptible and resistant *E*.*coli* strains were chosen to test the performance of novel AMR test. Kanamycin resistant (ΔtolC-KanR) and susceptible (D1) *E*.*coli* strains were compared (Figure 5A). The intensities of two formazan bands were followed in both samples varying concentration of kanamycin.

**Figure 5.**
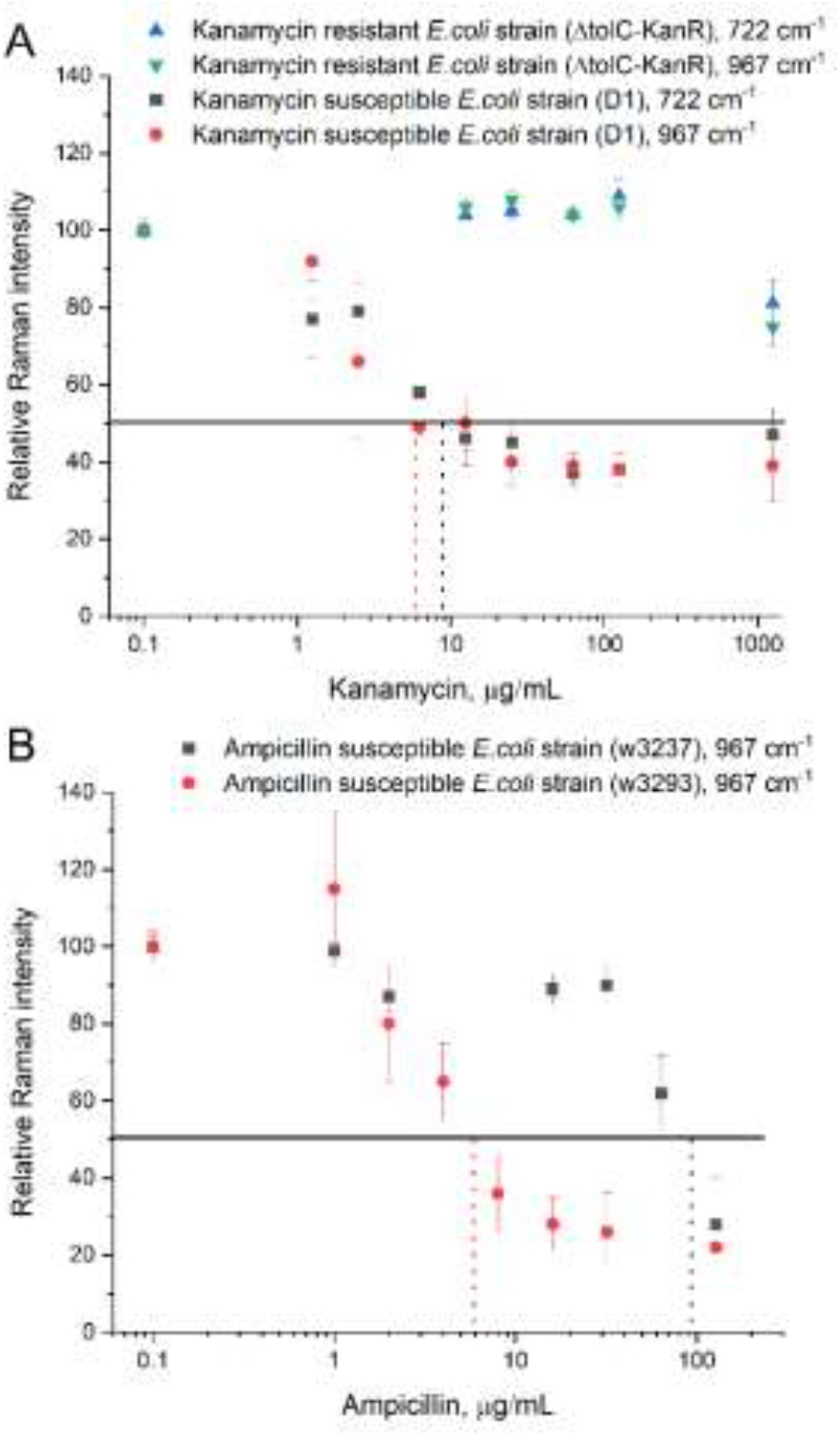
AMR test on the kanamycin susceptible and resistant E.coli strains (A) and ampicillin susceptible and resistant E.coli strains (B). 50% of Raman intensity was used as a reference value for antibiotic susceptibility.

The Raman intensities were normalized to the values from the sample without the antibiotic. The most intensive Raman bands of formazan, 722 and 967 cm^−1^, were compared. Both bands provided similar concentration dependencies (Figure 5A), so any of this bands can be used in the AMR test. MIC can be estimated taking 50% of Raman intensity as a reference value of susceptibility to the antibiotic. The MICs were 6 and 9 µg/mL for 967 and 722 cm^−1^ dependencies, respectively, for the kanamycin susceptible *E*.*coli* strain (D1). The kanamycin resistant *E*.*coli* strain (ΔtolC-KanR) had MIC larger than 1000 µg/mL.

Similar experiment was conducted with ampicillin resistant (w3237) and ampicillin susceptible (w3239) clinical strains of *E. coli* (Figure 5B). The 967 cm^−1^ band was chosen for evaluation. Again, at some concentration signal decreases rapidly, and further ampicillin concentration increase affects intensity slightly. Both strains had ampicillin dependent decrease in Raman signal, but MICs differ 15 times, namely 6 µg/mL for susceptible strain and 90 µg/mL of resistant strain.

### Direct comparison with conventional techniques in MIC determination

For this experiment, several clinical strains of *E. coli* and *K. pneumoniae* were chosen that are resistant or susceptible to levofloxacin. Minimal inhibitory concentrations of levofloxacin for these strains were established with a conventional AMR technique (Etest).

Simultaneously, MICs were determined with Raman resonance spectroscopy (a novel AMR test) using 50% decrease in Raman signal as a reference value. The results are provided in Table 1. The MICs determined with these two techniques were the same demonstrating good performance of the novel test.

**Table 1.**
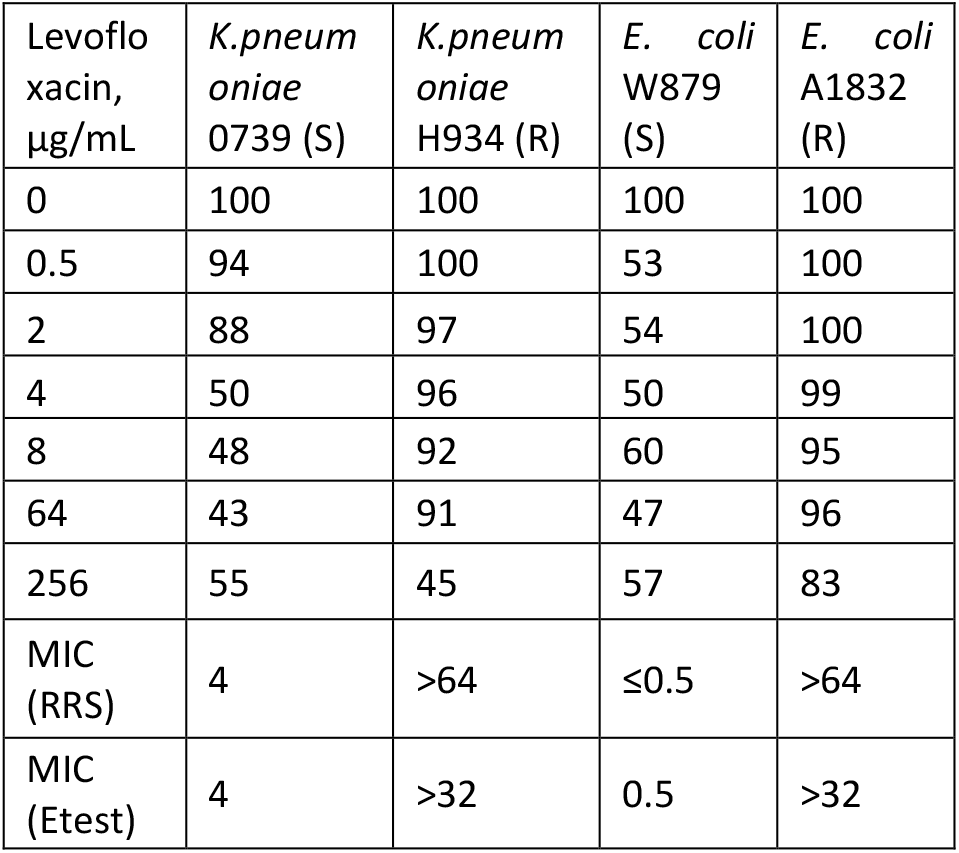
Comparison of MICs to levofloxacin determined with the novel AMR test (resonance Raman spectroscopy, RRS) and standard AMR (Etest). (S) means susceptible strain, (R) means resistant strain. The relative standard deviations varied from 5 to 10%.

| Levofloxacin, $\mu\text{g/mL}$ | *K.pneumoniae* 0739 (S) | *K.pneumoniae* H934 (R) | *E. coli* W879 (S) | *E. coli* A1832 (R) |
| --- | --- | --- | --- | --- |
| 0 | 100 | 100 | 100 | 100 |
| 0.5 | 94 | 100 | 53 | 100 |
| 2 | 88 | 97 | 54 | 100 |
| 4 | 50 | 96 | 50 | 99 |
| 8 | 48 | 92 | 60 | 95 |
| 64 | 43 | 91 | 47 | 96 |
| 256 | 55 | 45 | 57 | 83 |
| MIC (RRS) | 4 | >64 | $\leq 0.5$ | >64 |
| MIC (Etest) | 4 | >32 | 0.5 | >32 |

The developed AMR test provided a rapid determination of MIC during 1.5 hour for a fresh bacterial culture. The MTT test is known to be applicable for a wide variety of bacteria. Resonance Raman spectroscopy provides a possibility of independent specific determination of formazan without destroying of the cells with detergents. The method provided reliable MIC values for several strains of *E*.*coli* and *K*.*pneumoniae* and three types of antibiotics, including ampicillin from penicillin family, kanamycin from aminoglycoside family and levofloxacin from fluoroquinolone family.

## Materials and methods

### Bacterial cultures and reagents

Strains of *E*.*coli* (ΔtolC, D1, ΔtolC-KanR ^47,48^) were obtained from own collection of the laboratory of Olga Dontsova. Clinical samples (*K*.*pneumoniae* 0739, *K*.*pneumoniae* H934, *E. coli* W879, *E. coli* A1832, *E. coli* w3237, *E. coli* w3239) were obtained from Kulakov Medical Research Center, Moscow.

MTT bromine salt was obtained from Dia-M (Russia). MTT was dissolved in water, diluted to concentration of 4 mg/ml and stored at 4°C for a week. LB broth (Lennox) was purchased from Condalab (Italy). Antibiotics (levofloxacin, kanamycin, ampicillin) were purchased from Sigma-Aldrich (U.S.). Phosphate buffer solution (PBS) with pH 7.2 contained 0.8% NaCl, 0.02% KH_2_PO_4_, 0.02% KCl, and 0.12% Na_2_HPO_4_·12H_2_O. In this study MilliQ water was used. All chemicals were analytical grade reagents.

LB liquid medium was prepared from LB broth (Lennox) via manufacturer’s protocol. 4 g of the solid medium was suspended in 200mL of milliQ water, mixed and sterilized in autoclave for 15 minutes at 121°C. Prepared medium was stored at 4°C.

### Instruments and measurement parameters

In this study two Raman spectrometers were used. Spectrometers were bought from Enspectr (Russia) and Photon-Bio (Russia). 532nm wavelength Raman spectra were obtained on the Enspectr RaPort spectrometer, and 637 nm spectra were obtained on the Photon-Bio RL637 spectrometer.

Optical density measurements were made on ECROS 5400UV spectrophotometer (Russia).

Raman spectra for both wavelengths were collected with exposure time set at 1300 ms for 20 repeats. Laser powers for 532 nm and 637 nm excitation were set at 30mW and 100mW respectively.

Collected spectra were smoothed by Whittaker filtering algorithm^49^ and baselines were corrected with asymmetrically reweighted penalized least squares smoothing.^50^

### Raman – assisted MTT antibiotic resistance assay

Bacterial strains were obtained from overnight cultures, diluted with media to 2 MFU (McFarland Units, around 6^*^10^8^ cells/ml, cell concentration was measured on spectrophotometer at 570 nm, OD_570_ for 2MFU = 0.451) and incubated at 37°C for an hour. Then 50 μL of cell culture was added to 350 μL of PBS with the required amount of antibiotic and incubated at 37°C for another 1h. After the incubation, 100 μL MTT was added to the samples, and after the incubation for 20 minutes, samples (500 μL each) were placed into glass vials and measured with a spectrometer.

Intensity of formazan peak at 967 cm^−1^ were collected from each spectrum from samples with antibiotic and related to those of the control sample without antibiotic. The final intensity values were presented as a percentage of the intensity of the control sample.

## References

1. Murray, Christopher JL, et al. “Global burden of bacterial antimicrobial resistance in 2019: a systematic analysis.” The Lancet 399.10325 (2022): 629–655

2. Durand, Guillaume André, Didier Raoult, and Grégory Dubourg. “Antibiotic discovery: history, methods and perspectives.” International journal of antimicrobial agents 53.4 (2019): 371–382.

3. Dadgostar, Porooshat. “Antimicrobial resistance: implications and costs.” Infection and drug resistance (2019): 3903–3910.

4. Yang, Chien-Chang, et al. “Risk factors of treatment failure and 30-day mortality in patients with bacteremia due to MRSA with reduced vancomycin susceptibility.” Scientific Reports 8.1 (2018): 7868.

5. Burnham, Carey-Ann D., et al. “Diagnosing antimicrobial resistance.” Nature Reviews Microbiology 15.11 (2017): 697–703.

6. Bonine, Nicole Gidaya, et al. “Impact of delayed appropriate antibiotic therapy on patient outcomes by antibiotic resistance status from serious gramnegative bacterial infections.” The American journal of the medical sciences 357.2 (2019): 103–110.

7. Ghasemi, Mahshid, et al. “The MTT assay: utility, limitations, pitfalls, and interpretation in bulk and single-cell analysis.” International journal of molecular sciences 22.23 (2021): 12827.

8. Grela, Ewa, Joanna Kozłowska, and Agnieszka Grabowiecka. “Current methodology of MTT assay in bacteria– A review.” Acta histochemica 120.4 (2018): 303–311.

9. Foongladda, S., et al. “Rapid and simple MTT method for rifampicin and isoniazid susceptibility testing of Mycobacterium tuberculosis.” The International Journal of Tuberculosis and Lung Disease 6.12 (2002): 1118–1122.

10. Riss, Terry. “Is your MTT assay really the best choice.” Promega™, Madison, Wisconsin, United States) (2017).

11. Mao, Zhu, et al. “Predictive value of the surface-enhanced resonance Raman scattering-based MTT assay: a rapid and ultrasensitive method for cell viability in situ.” Analytical chemistry 85.15 (2013): 7361–7368.

12. Sharma, Bhavya, et al. “SERS: Materials, applications, and the future.” Materials today 15.1-2 (2012): 16–25.

13. Efremov, Evtim V., Freek Ariese, and Cees Gooijer. “Achievements in resonance Raman spectroscopy: Review of a technique with a distinct analytical chemistry potential.” Analytica chimica acta 606.2 (2008): 119–134.

14. Mao, Zhu, et al. “In situ semi-quantitative assessment of single-cell viability by resonance Raman spectroscopy.” Chemical Communications 54.52 (2018): 7135–7138.

15. Van Meerloo, Johan, Gertjan JL Kaspers, and Jacqueline Cloos. “Cell sensitivity assays: the MTT assay.” Cancer cell culture: methods and protocols (2011): 237–245.

16. Montoro, Ernesto, et al. “Comparative evaluation of the nitrate reduction assay, the MTT test, and the resazurin microtitre assay for drug susceptibility testing of clinical isolates of Mycobacterium tuberculosis.” Journal of Antimicrobial Chemotherapy 55.4 (2005): 500–505.

17. Mshana, Robert N., et al. “Use of 3-(4, 5-dimethylthiazol-2-yl)-2, 5-diphenyl tetrazolium bromide for rapid detection of rifampin-resistant Mycobacterium tuberculosis.” Journal of clinical microbiology 36.5 (1998): 1214–1219.

18. Brambilla, Eugenio, et al. “The influence of antibacterial toothpastes on in vitro Streptococcus mutans biofilm formation: a continuous culture study.” Am J Dent 27.3 (2014): 160–6.

19. Stevens, Mark G., Marcus E. Kehrli Jr, and Peter C. Canning. “A colorimetric assay for quantitating bovine neutrophil bactericidal activity.” Veterinary immunology and immunopathology 28.1 (1991): 45–56.

20. English, B. Keith, and Aditya H. Gaur. “The use and abuse of antibiotics and the development of antibiotic resistance.” Hot topics in infection and immunity in children VI (2010): 73–82.

21. Jagusztyn-Krynicka, Elbieta K., and Agnieszka Wyszynska. “The decline of antibiotic era–new approaches for antibacterial drug discovery.” Polish Journal of Microbiology 57.2 (2008): 91.

22. Dickinson, John D., and Marin H. Kollef. “Early and adequate antibiotic therapy in the treatment of severe sepsis and septic shock.” Current infectious disease reports 13 (2011): 399–405.

23. Syal, Karan, et al. “Current and emerging techniques for antibiotic susceptibility tests.” Theranostics 7.7 (2017): 1795.

24. Paul, Mical, et al. “Systematic review and meta-analysis of the efficacy of appropriate empiric antibiotic therapy for sepsis.” Antimicrobial agents and chemotherapy 54.11 (2010): 4851–4863.

25. Tolosa, Laia, María Teresa Donato, and María José Gómez-Lechón. “General cytotoxicity assessment by means of the MTT assay.” Protocols in in vitro hepatocyte research (2015): 333–348.

26. Weichert, H., et al. “The MTT-assay as a rapid test for cell proliferation and cell killing: application to human peripheral blood lymphocytes (PBL).” Allergie und Immunologie 37.3-4 (1991): 139–144.

27. Adan, Aysun, Yagmur Kiraz, and Yusuf Baran. “Cell proliferation and cytotoxicity assays.” Current pharmaceutical biotechnology 17.14 (2016): 1213–1221.

28. Molaae, Nada, et al. “Evaluating the proliferation of human PeripheralBlood mononuclear cells using MTT assay.” International Journal of Basic Science in Medicine 2.1 (2017): 25–28.

29. Yuan, Yuan, et al. “In vitro screening of five Hainan plants of Polyalthia (Annonaceae) against human cancer cell lines with MTT assay.” J Med Plants Res 5 (2011): 837–841.

30. Nafie, Mohamed S., Mohamed A. Tantawy, and Gamal A. Elmgeed. “Screening of different drug design tools to predict the mode of action of steroidal derivatives as anti-cancer agents.” Steroids 152 (2019): 108485.

31. Supino, Rosa. “MTT assays.” In vitro toxicity testing protocols (1995): 137–149.

32. Dodo, Kosuke, Katsumasa Fujita, and Mikiko Sodeoka. “Raman spectroscopy for chemical biology research.” Journal of the American Chemical Society 144.43 (2022): 19651–19667.

33. Pezzotti, Giuseppe. “Raman spectroscopy in cell biology and microbiology.” Journal of Raman Spectroscopy 52.12 (2021): 2348–2443.

34. Petry, Renate, Michael Schmitt, and Jürgen Popp. “Raman spectroscopy—a prospective tool in the life sciences.” chemphyschem 4.1 (2003): 14-30.

35. Robert, Bruno. “Resonance raman spectroscopy.” Photosynthesis research 101 (2009): 147–155.

36. Efremov, Evtim V., et al. “Strong overtones and combination bands in ultraviolet resonance Raman spectroscopy.” Analytical chemistry 78.9 (2006): 3152–3157.

37. Moskovits, Martin, et al. “SERS and the single molecule.” Optical properties of nanostructured random media. Berlin, Heidelberg: Springer Berlin Heidelberg, 2002. 215–227.

38. Mosier-Boss, Pamela A. “Review of SERS substrates for chemical sensing.” Nanomaterials 7.6 (2017): 142.

39. Grys, David-Benjamin, et al. “Eliminating irreproducibility in SERS substrates.” Journal of Raman Spectroscopy 52.2 (2021): 412–419.

40. Moodley, Suventha, et al. “The 3-(4, 5-dimethylthiazol-2-yl)-2, 5-diphenyl tetrazolium bromide (MTT) assay is a rapid, cheap, screening test for the in vitro anti-tuberculous activity of chalcones.” Journal of microbiological methods 104 (2014): 72–78.

41. Meyers, Annika, Christoph Furtmann and Joachim Jose. “Direct optical density determination of bacterial cultures in microplates for high-throughput screening applications”. Enzyme and Microbial Technology 118 (2018): 1–5.

42. Mira, Portia, Pamela Yeh and Barry G. Hall. “Estimating microbial population data from optical density”. PLoS ONE 17(10): e0276040.

43. Shi, Lei, et al. “Synthesis and antimicrobial activities of Schiff bases derived from 5-chloro-salicylaldehyde.” European journal of medicinal chemistry 42.4 (2007): 558–564.

44. Benov, Ludmil. “Effect of growth media on the MTT colorimetric assay in bacteria.” PloS one 14.8 (2019): e0219713.

45. Chakrabarti, Ranjana, et al. “Vitamin A as an enzyme that catalyzes the reduction of MTT to formazan by vitamin C.” Journal of Cellular Biochemistry 80.1 (2001): 133–138.

46. Han, Mei, et al. “Limitations of the use of MTT assay for screening in drug discovery.” J Chin Pharmaceu Sci 19 (2010): 195–200.

47. Baba T., et al. “Construction of Escherichia coli K-12 in-frame, singlegene knockout mutants: the Keio collection”. Mol. Syst. Biol. 2006; 2: 2006.0008.

48. Mehla K, Ramana J. Structural signature of Ser83Leu and Asp87Asn mutations in DNA gyrase from enterotoxigenic Escherichia coli and impact on quinolone resistance. Gene. 2016; 576(1 Pt 1):28–35.

49. Eilers P. H. C. A perfect smoother //Analytical chemistry. –2003. – ?. 75. – ?. 14. – ?. 3631–3636.

50. Baek, Sung-June, et al. “Baseline correction using asymmetrically reweighted penalized least squares smoothing.” Analyst 140.1 (2015): 250–257.

